# Decoupling of CO_2_ and O_2_ fluxes reveals oxygen-independent decarboxylation in anoxic mammalian cells

**DOI:** 10.64898/2026.10.10.757797

**Authors:** Hans Vellama, Hillar Eichelmann, Uku Gross, Hendrik Luuk, Kattri-Liis Eskla

## Abstract

Canonical understanding of cellular respiration holds that CO_2_ production tracks oxygen consumption, and mammalian cells deprived of oxygen are thought to rely on lactate fermentation, which yields no net CO_2_. Because standard respirometry infers CO_2_ flux either from oxygen consumption or from extracellular acidification rather than measuring it directly, a CO_2_ flux decoupled from oxygen would go undetected or be misattributed to glycolysis. Using direct measurement of CO_2_ and O_2_ fluxes from the gas phase in anoxic mammalian cells, we show that a substantial CO_2_ efflux persists after oxygen is fully removed. This anoxic flux scales linearly with normoxic CO_2_ and with cell mass, and exhibits a lower activation energy than normoxic respiration, suggesting a distinct rate-limiting chemistry. Pyruvate and α-ketobutyrate increased the anoxic CO_2_ flux while lactate dehydrogenase inhibition abolished the pyruvate-induced increase, supporting a role for cytosolic NAD^+^ regeneration in oxygen-independent decarboxylation. ^13^C tracing identified a substantial glutamine-derived component, consistent with mitochondrial decarboxylation, and a smaller glucose-derived component associated with the cytosolic pentose phosphate pathway. Direct CO_2_ measurement thus reframes lactate dehydrogenase as an enabler of oxygen-independent decarboxylation.

**SYNOPSIS:** Direct measurement of CO_2_ and O_2_ fluxes reveals that mammalian cells continue to release CO_2_ after oxygen consumption ceases. This oxygen-independent CO_2_ production comprises a major glutamine-derived component and a smaller glucose-derived component associated with the pentose phosphate pathway, with the latter constrained by cytosolic NAD^+^ regeneration through lactate dehydrogenase.

## INTRODUCTION

In aerobic metabolism, the mitochondrial electron transport chain reduces oxygen to water while the decarboxylation reactions of the tricarboxylic acid (TCA) cycle release CO_2_. Most methods for measuring respiration rely on the assumption that TCA activity (and thus CO_2_ production) follows oxygen consumption, because these decarboxylation and oxidation reactions are linked through the NAD^+^/NADH couple (Kory *et al*, 2020). High-resolution respirometry measures only oxygen consumption while extracellular flux analysis estimates the respiratory contribution to acidification from the oxygen consumption rate rather than from CO_2_ itself (Mookerjee *et al*, 2015). Extracellular acidification also reflects two proton sources, lactate export and the hydration of respiratory CO_2_, and cannot separate them without further assumptions. A CO_2_ flux that does not follow oxygen consumption would therefore be missed or assigned to glycolysis. Direct measurement of CO_2_ release from mammalian cells is uncommon, and the sources of CO_2_ flux once oxygen consumption approaches zero have not been examined directly. Under anoxia, mammalian cells are generally expected to increase anaerobic glycolysis where glucose is converted to pyruvate that is further reduced to lactate (Grimm *et al*, 2024). The latter step regenerates NAD^+^ from NADH but produces no net CO_2_. Unlike yeast, mammalian cells lack pyruvate decarboxylase, which converts pyruvate to acetaldehyde and CO_2_ during alcoholic fermentation. The oxidative decarboxylations that account for most CO_2_ output under normoxia, catalyzed by pyruvate dehydrogenase, isocitrate dehydrogenase and α-ketoglutarate dehydrogenase, all use NAD^+^ as an electron acceptor (Mookerjee *et al*, 2015; Arnold & Finley, 2023). In the conventional view, the bulk of NAD^+^ is regenerated by the oxygen-dependent electron transport chain, implying that anoxia is expected to stop these reactions and remove the main source of respiratory CO_2_ (Chinopoulos, 2013).

Here, we measure CO_2_ and O_2_ fluxes simultaneously and independently in the gas phase above adherent cells, enabling direct calculation of the respiratory quotient (RQ) when O_2_ consumption is measurable and assessment of CO_2_ production independently of oxygen consumption throughout normoxia–anoxia–reoxygenation transitions. We find that a substantial CO_2_ efflux continues after oxygen consumption has stopped. This efflux scales with cell mass and comprises a substantial glutamine-derived component, consistent with mitochondrial decarboxylation, and a smaller glucose-derived component associated with the cytosolic pentose phosphate pathway. Our findings further implicate lactate dehydrogenase-dependent NAD^+^ regeneration in regulating anoxic CO_2_ production. Direct measurement of CO_2_ thus reveals an oxygen-independent decarboxylation that respirometry based on oxygen consumption would not detect.

## RESULTS AND DISCUSSION

### CO_2_ efflux persists in anoxic cells

To measure respiratory CO_2_ directly rather than infer it from oxygen consumption or medium acidification, we monitored CO_2_ and O_2_ concentration changes simultaneously in the gas phase above cultured HKC8 and HepG2 cells (Fig 1A). Lowering gas-phase O_2_ to anoxia (O_2_ <0.03%) abolished O_2_ flux, yet a substantial CO_2_ efflux persisted and stabilized at a non-zero plateau, representing ∼10% and ∼15% of the corresponding normoxic CO_2_ flux in HKC8 and HepG2 cells, respectively (Fig 1A-B). Restoring oxygen returned both fluxes toward their normoxic values, indicating that the anoxic efflux is a reversible metabolic output rather than a consequence of cell death or membrane leak (Fig 1A). Because CO_2_ and O_2_ fluxes were measured independently, the respiratory quotient (RQ = CO_2_/O_2_) could be calculated directly from the measured fluxes when O_2_ consumption was nonzero (Fig 1A). Under anoxia, RQ is undefined because O_2_ consumption ceases. In both cell lines, supplementation with pyruvate significantly increased anoxic CO_2_ production. Inhibition of the malate-aspartate shuttle (MAS), which transfers reducing equivalents between cytosol and mitochondria, with aminooxyacetic acid (AOAA) did not significantly affect anoxic CO_2_ production in HKC8 cells but reduced it in HepG2 cells (Fig 1B).

**Figure 1.**
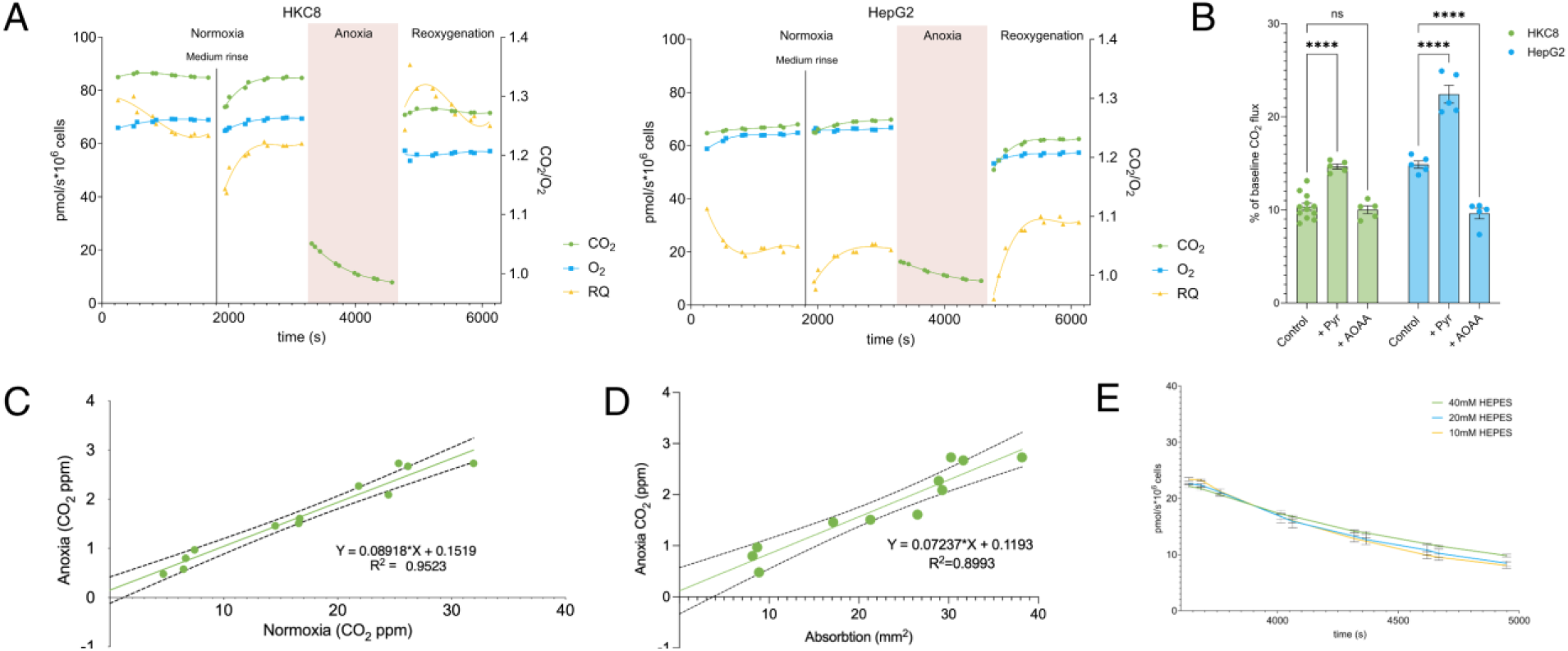
CO_2_ efflux persists in anoxic cells. (A) Time course of simultaneously measured O_2_ consumption and CO_2_ production during a normoxia–anoxia–reoxygenation cycle in HKC8 and HepG2 cells. RQ was calculated as CO_2_/O_2_ where O_2_ consumption was nonzero; RQ is undefined under anoxia. Cells were rinsed with their initial medium during normoxia, as indicated. Data represent the mean of three independent biological experiments. **(B)** Anoxic CO_2_ flux, expressed as a percentage of the corresponding normoxic baseline CO_2_ flux, in HKC8 and HepG2 cells incubated in complete respiration medium or supplemented with pyruvate (3 mM) or with the transaminase/malate-aspartate shuttle inhibitor aminooxyacetic acid (AOAA, 5 mM). **(C)** Correlation between normoxic and anoxic CO_2_ production in HKC8 cells. Solid line, linear regression; dashed lines, 95% confidence interval. **(D)** Correlation between cell mass estimated by optical absorption and anoxic CO_2_ production in HKC8 cells. Solid line, linear regression; dashed lines, 95% confidence interval. (E) Time course of anoxic CO_2_ efflux in HKC8 cells at different HEPES concentrations in the measurement medium (10, 20, and 40 mM). Data represent the mean of three independent experiments. (B, E) Data are presented as mean ± SEM. Each dot in panels B-D represents one independent biological experiment. Statistical analysis: (B) One-way ANOVA followed by Holm-Šídák’s multiple-comparisons test. (C,D) Simple linear regression. \*\*\*\**P* < 0.0001.

### Anoxic CO_2_ efflux scales with cell mass

The magnitude of the anoxic efflux scaled linearly with the normoxic CO_2_ rate (r^2^ = 0.95; Fig 1C) and with cell number measured by optical absorbance (r^2^ = 0.90; Fig 1D), indicating that it tracks the metabolic capacity and mass of the cell population rather than a methodological source. Although HEPES was the primary buffer in the respiration medium, a small bicarbonate-buffering component remains due to the ∼550 ppm CO_2_ supplied in the gas mixture. To assess whether medium acidification could contribute substantially to the anoxic CO_2_ signal, CO_2_ efflux was measured at different HEPES concentrations (Fig 1E). If buffer acidification were a major source of the measured CO_2_, altering the buffering capacity would be expected to affect CO_2_ efflux. Theoretical calculations predicted a greater contribution from bicarbonate loss at the lowest HEPES concentration (Table EV1). However, despite differences in pH between the buffering conditions, the measured anoxic CO_2_ efflux remained similar across all three concentrations (Fig 1E), arguing against medium acidification as the predominant source of the signal and supporting a substantial cellular contribution.

### Anoxic CO_2_ is produced by a distinct decarboxylating chemistry

The anoxic efflux correlated only weakly with mitochondrial DNA content (r^2^ = 0.48, Fig 2A) and persisted, at reduced levels, in HepG2 cells depleted of mitochondrial DNA by ethidium bromide treatment (Fig 2B). Consistent with this, acute inhibition of mitochondrial complex I with rotenone did not eliminate CO_2_ production. In both HepG2 and HKC8 cells, substantial CO_2_ production persisted in the presence of rotenone under normoxia and was further reduced, but remained detectable, when rotenone-treated cells were subsequently exposed to anoxia (Fig 2C). Thus, some anoxic CO_2_ production persists when mitochondrial electron transport is restricted by either chronic mtDNA depletion or acute complex I inhibition. To further assess whether anoxic CO_2_ production reflects a process distinct from normoxic respiration, we examined its temperature dependence. The apparent activation energy was 24 kJ·mol^−1^ under anoxia against 65 kJ·mol^−1^ in normoxia (P < 0.0001; n = 23 experiments in total) (Fig 2D), indicating a different rate-limiting step.

**Figure 2.**
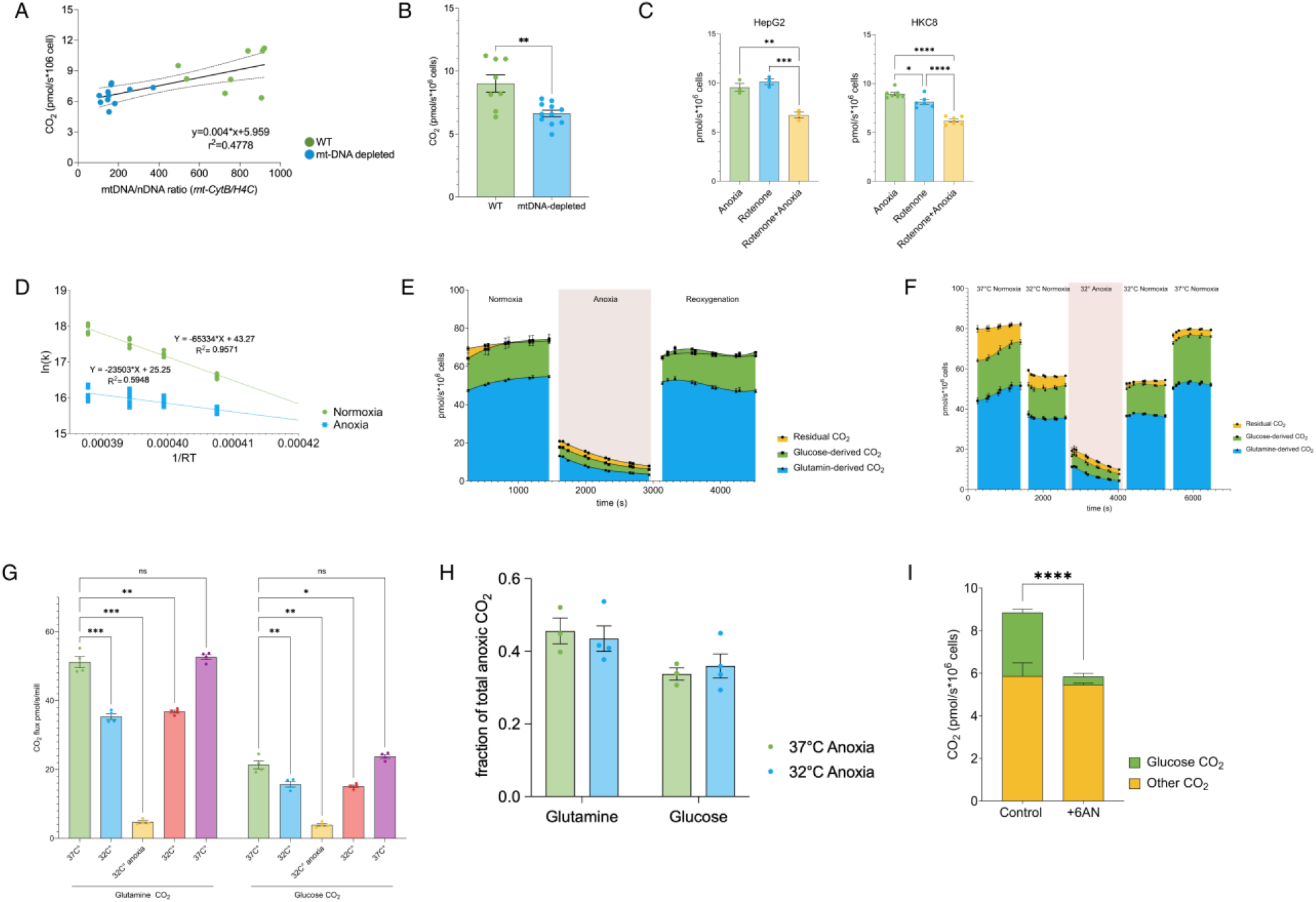
Anoxic CO_2_ is produced by a distinct decarboxylating chemistry. (A) Correlation between relative mitochondrial DNA content, expressed as the mtDNA/nDNA ratio (*mt-CytB/H4C*), and anoxic CO_2_ production. Green symbols represent wild-type (WT) HepG2 cells and blue symbols represent mtDNA-depleted HepG2 cells. Solid line, linear regression; dashed lines, 95% confidence interval. **(B)** Anoxic CO_2_ production in WT and mtDNA-depleted HepG2 cells. **(C)** CO_2_ production in HepG2 and HKC8 cells during anoxia, during inhibition of mitochondrial complex I with rotenone (0.5 μM) under normoxia, or during rotenone treatment followed by anoxia. **(D)** Arrhenius plot of CO_2_ production in HKC8 cells under normoxic and anoxic conditions. The natural logarithm of CO_2_ flux (k) was plotted against 1/RT, where R is the universal gas constant and T is the absolute temperature. Apparent activation energies (E_a_) were calculated as the negative slope of the linear regression: 65 kJ·mol^−1^ under normoxia and 24 kJ·mol^−1^ under anoxia. **(E)** ^13^C-tracing analysis of CO_2_ production during a normoxia–anoxia–reoxygenation cycle in HKC8 cells. Glucose- and glutamine-derived CO_2_ fluxes were estimated separately using uniformly ^13^C-labelled glucose or glutamine. Stacked areas show the contributions of glutamine-derived, glucose-derived, and residual CO_2_ to total CO_2_ production. Residual CO_2_ represents the remaining CO_2_ flux after subtracting the estimated glucose- and glutamine-derived contributions from the total measured CO_2_ flux. **(F)** ^13^C-tracing analysis of CO_2_ production in HKC8 cells during normoxia at 37 °C, cooling to 32 °C, anoxia and reoxygenation at 32 °C, and subsequent rewarming to 37 °C. Stacked areas show glutamine-derived, glucose-derived, and residual CO_2_ fluxes, as defined in (E). **(G)** Absolute glutamine- and glucose-derived CO_2_ fluxes quantified from the ^13^C-tracing experiment in (F) at each experimental stage. **(H)** Relative contributions of glutamine- and glucose-derived CO_2_ to total anoxic CO_2_ production in HKC8 cells at 37 °C and 32 °C, expressed as fractions of the total anoxic CO_2_ flux. **(I)** ^13^C-tracing analysis of anoxic CO_2_ production following pretreatment with the pentose phosphate pathway inhibitor 6-aminonicotinamide (6-AN, 300 μM, 16 h) in HKC8 cells. Statistical comparison was performed between the glucose-derived CO_2_ fractions in complete and 6-AN-treated conditions. Inhibition of the pentose phosphate pathway selectively reduced the glucose-derived CO_2_ fraction, whereas the remaining CO_2_ fraction (including glutamine-derived CO_2_ and residual CO_2_) was largely unchanged. Data are presented as mean ± SEM. Each dot in panels A-D, G-H represents one independent biological experiment. Panels E, F and I show the mean contribution to CO_2_ production calculated from 3-5 independent biological experiments. Statistical analysis: (A,D) simple linear regression; (B, I) two-tailed Mann-Whitney test; (C) one-way ANOVA followed by Tukey’s multiple-comparisons test; (G) two-way ANOVA followed by Dunnett’s multiple comparisons test. \**P* < 0.05, \*\**P* < 0.01, \*\*\**P* < 0.001, \*\*\*\**P* < 0.0001.

To identify the relative contribution of glucose and glutamine to CO_2_ production in the complete medium, cells were co-incubated with glucose and glutamine and each substrate was replaced in turn by its uniformly ^13^C-labelled form. Since it is well-established that the LI-COR CO_2_ analyzer under-detects ^13^CO_2_ by 69% relative to CO_2_ (McDermitt *et al,* 1995), the reduction in the measured flux corresponds to the CO_2_ derived from the labeled substrate, enabling the estimation of the corrected CO_2_ contribution of the labeled substrate under the assumption of identical total CO_2_ production on unlabeled and labeled substrates. Across a normoxia-anoxia-reoxygenation cycle, total CO_2_ production declined markedly during anoxia and recovered upon reoxygenation. Despite the loss of oxidative metabolism, a substantial glucose-derived CO_2_ flux persisted throughout anoxia, contributing approximately 3.0 pmol·s^−1^·(10^6^ cells)^−1^ at the anoxic endpoint alongside 3.8 pmol·s^−1^·(10^6^ cells)^−1^ from glutamine and 2.0 pmol·s^−1^·(10^6^ cells)^−1^ as residual CO_2_ (Fig 2E).

We next asked whether hypothermia altered the magnitude or substrate composition of this anoxic CO_2_ production. Cooling HKC8 cells from 37 °C to 32 °C reduced both glucose- and glutamine-derived CO_2_ flux under normoxia, with a further pronounced decline in both components upon transition to anoxia at 32 °C (Fig 2F,G). Reoxygenation at 32 °C restored both substrate-derived fluxes, and subsequent return to 37 °C restored them to initial normothermic levels (Fig 2F,G). Despite these marked changes in absolute CO_2_ production, the fractional contributions of glucose and glutamine to total anoxic CO_2_ were not significantly different between 37 °C and 32 °C (Fig 2H). Thus, hypothermia suppresses the overall rate of anoxic carbon decarboxylation without detectably shifting the relative contribution of glucose versus glutamine to total anoxic CO_2_ production. To test the origin of the glucose-derived component more directly, inhibition of the pentose phosphate pathway using 6-AN significantly suppressed anoxic CO_2_ production, with the reduction occurring predominantly in the glucose-derived component (Fig 2I). Together, these results suggest that anoxic CO_2_ production includes contributions from multiple substrates and remains detectable despite impaired mitochondrial electron transport, while the glucose-derived component is consistent with pentose phosphate pathway-associated decarboxylation rather than TCA-cycle oxidation.

### Anoxic CO_2_ efflux is limited by cytosolic NAD^+^ regeneration through LDH

Pyruvate and α-ketobutyrate, each of which regenerates NAD^+^ through lactate dehydrogenase, increased the anoxic CO_2_ efflux (up to ∼17.5% of normoxic CO_2_ flux) (Fig 3A, B). The LDH inhibitor oxamate abolished this increase (Fig 3A). Inhibiting the transaminases and malate-aspartate shuttle (AOAA), succinate dehydrogenase (malonate), the citrate carrier (BCA), or the mitochondrial pyruvate carrier (UK5099) failed to abolish the pyruvate-induced increase (Fig 3A). The efflux is therefore limited by the supply of cytosolic NAD^+^ rather than by oxygen. This role places lactate dehydrogenase, the conventional endpoint of CO_2_-free fermentation, as an enabler of oxygen-independent decarboxylation. Consistent with a redox-limited mechanism, α-ketobutyrate selectively increased glucose-derived CO_2_ production during anoxia, indicating that glucose-derived decarboxylation is constrained by cytosolic NAD^+^ availability rather than carbon substrate availability (Fig 3C).

**Figure 3.**
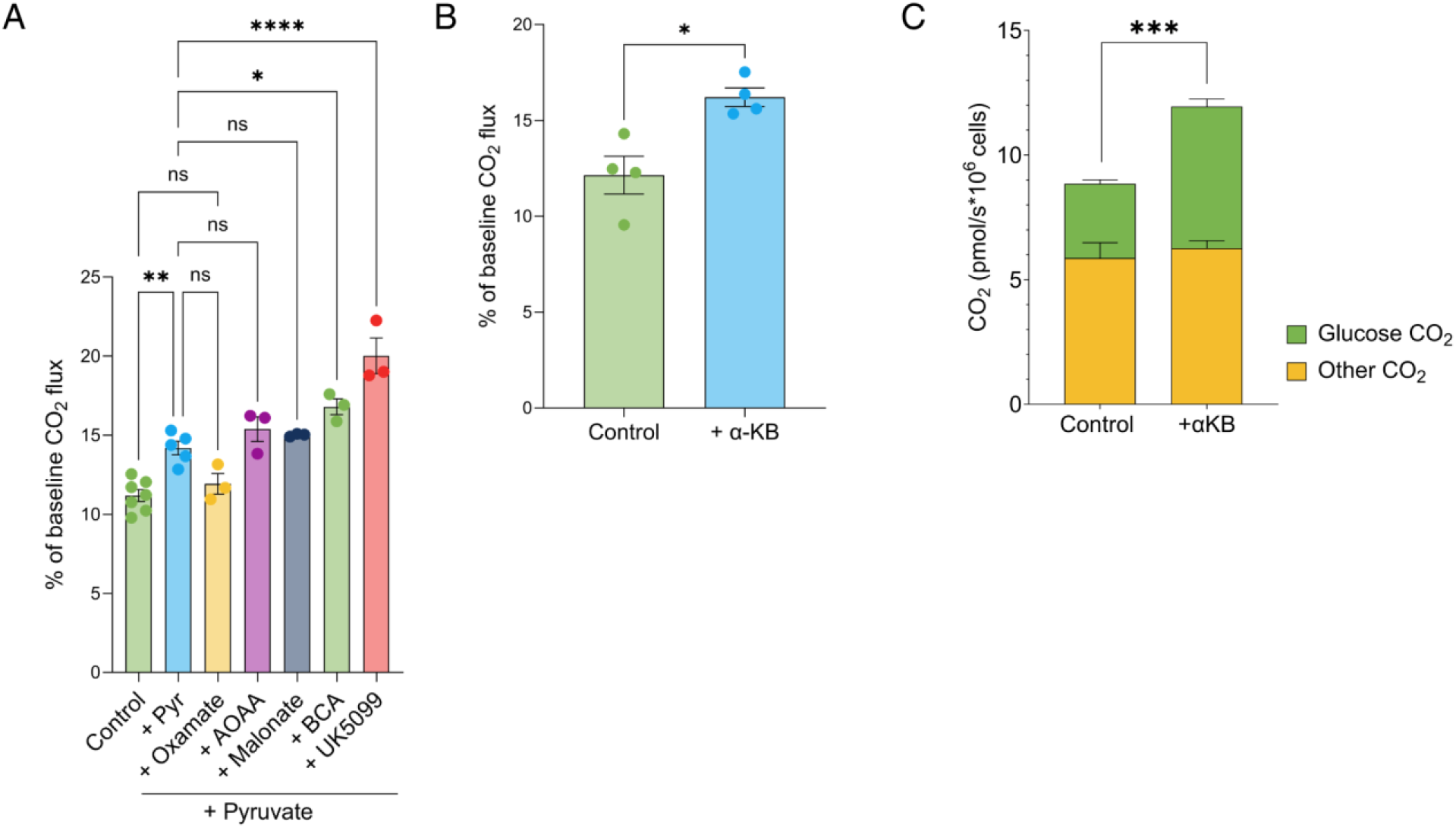
Anoxic CO_2_ efflux is limited by cytosolic NAD^+^ regeneration through LDH. (A) Anoxic CO_2_ flux, expressed as a percentage of the corresponding normoxic baseline CO_2_ flux, in HKC8 cells incubated in complete respiration medium or supplemented with pyruvate (3 mM), pyruvate (3 mM) plus the LDH inhibitor oxamate (100 mM), or pyruvate together with inhibitors of the malate-aspartate shuttle/transaminases AOAA (5 mM), succinate dehydrogenase inhibitor malonate (10 mM), the citrate carrier BCA (5 mM), or the mitochondrial pyruvate carrier UK5099 (4 μM). **(B)** α-Ketobutyrate (α-KB, 3 mM), an alternative electron acceptor that regenerates NAD^+^ through LDH, increased anoxic CO_2_ flux in HKC8 cells. **(C)** ^13^C-tracing analysis of anoxic CO_2_ production showing that α-KB (3 mM) selectively increased the glucose-derived CO_2_ fraction, whereas the remaining CO_2_ fraction (including glutamine-derived CO_2_ and residual CO_2_) was largely unchanged in HKC8 cells. Statistical comparison was performed between the glucose-derived CO_2_ fractions in complete and α-KB-treated conditions. Data are presented as mean ± SEM. Each dot in panels A and B represents one independent biological experiment. Panel C shows the mean contribution to CO_2_ production calculated from 3-4 independent biological experiments. Statistical analysis: (A) one-way ANOVA followed by Šídák’s multiple-comparisons test; (B,C) two-tailed unpaired Welch’s t-test. \**P* < 0.05, \*\**P* < 0.01, \*\*\**P* < 0.001, \*\*\*\**P* < 0.0001, ns - not significant.

Respiration is canonically viewed as a well-coordinated process in which CO_2_ production follows oxygen consumption. Direct measurement of the two fluxes shows that this coupling is conditional on the metabolic state of the cell. Even in normoxia, decarboxylation slightly exceeds oxygen consumption, presumably because CO_2_ is produced by multiple decarboxylases whereas oxygen is consumed mainly at cytochrome c oxidase (Fig 1A). Oxygen deprivation is just one instance of the more general electron-acceptor supply limitation, which slows nutrient oxidation without halting decarboxylation as indicated by persistent anoxic CO_2_ efflux.

The selective increase in glucose-derived anoxic CO_2_ following α-ketobutyrate treatment suggests that this component is sensitive to cellular redox state, whereas the glutamine-derived component is comparatively unaffected (Fig 3C). As an alternative electron acceptor, α-ketobutyrate can facilitate NADH oxidation and NAD^+^ regeneration through lactate dehydrogenase, potentially relieving constraints imposed by limited electron-acceptor availability. This is consistent with the importance of NAD^+^ regeneration for sustaining metabolism when mitochondrial electron transport is impaired (Sullivan *et al*, 2015; Titov *et al*, 2016). However, the glucose-derived CO_2_ is associated with the oxidative pentose phosphate pathway (Fig 2I and 3C), whose dehydrogenases require NADP^+^ rather than NAD^+^. The effect of α-ketobutyrate on this pathway is therefore likely indirect. Although NAD^+^ provides the substrate for NADP^+^ synthesis by NAD kinase (Hoxhaj *et al*, 2019), whether altered NAD^+^ availability influences NADP^+^ availability or pentose phosphate pathway activity under anoxia remains unresolved.

In the present study, glutamine was the largest identified contributor to anoxic CO_2_ production, exceeding the glucose-derived contribution at the anoxic endpoint (Fig 2E). Decarboxylation of α-ketoglutarate by α-ketoglutarate dehydrogenase reduces NAD^+^ to NADH in the mitochondrial matrix, so sustained flux under anoxia requires an oxygen-independent route for mitochondrial NADH reoxidation. The malate–aspartate shuttle is unlikely to be the main route, since aminooxyacetate, which blocks its aminotransferase steps, did not significantly affect anoxic CO_2_ flux in HKC8 cells (Fig 1B). An alternative is residual Complex I activity, sustained by Complex II operating in reverse as a fumarate reductase that reoxidises ubiquinol and thereby provides oxidized ubiquinone as the electron acceptor for Complex I (Spinelli *et al*, 2021; Ravasz *et al*, 2024). The reduction, but not elimination, of anoxic CO_2_ production by rotenone and by mtDNA depletion (Fig 2B, C) is consistent with a partial contribution of this route, while the remaining flux suggests that additional mechanisms are involved.

Methods that infer CO_2_ from oxygen consumption, or from extracellular acidification, return values close to zero for the anoxic decarboxylation flux and either miss it or assign it to glycolysis. This may be one reason why oxygen-independent decarboxylation has received relatively little attention. Measuring CO_2_ directly is expected to reveal additional features of energy metabolism that oxygen-based readouts cannot resolve, particularly in tissues with distinctive redox and substrate handling, such as adipose, muscle and neural tissue. The same reasoning applies to pathological states such as ischemia and severe hypoxia, where redox imbalance is a defining feature. Whether the decarboxylation described here is a general property of mammalian cells or is particularly prominent in metabolically active epithelial cell types such as HKC8 and HepG2 will require experiments across a wider range of cell types.

## METHODS

### Cell lines

Human kidney proximal tubular epithelial (HKC8) cells, kindly provided by Professor Sir Peter J. Ratcliffe, and HepG2 cells were cultured in low-glucose Minimum Essential Medium (Capricorn Scientific, Cat. No. MEM-STA) supplemented with 10% fetal bovine serum (Gibco, Thermo Fisher Scientific, Cat. No. A5256701) and 1× penicillin/streptomycin (Capricorn Scientific, Cat. No. PS-B) at 37 °C in a humidified atmosphere containing 5% CO_2_. Unless otherwise stated, all experiments were performed using HKC8 cells.

### Reagents

Metabolic substrates used in this study included sodium pyruvate (Corning, Cat. No. 25-000-CI) and α-ketobutyric acid (α-KB, Sigma-Aldrich, Cat. No. K401). Pharmacological inhibitors included sodium oxamate (Sigma-Aldrich, Cat. No. O2751), UK5099 (MedChemExpress, Cat. No. HY-15475), aminooxyacetic acid (AOAA; Sigma-Aldrich, Cat. No. C13408), malonate (Sigma-Aldrich, Cat. No. M1296-5G), 1,2,3-benzenetricarboxylic acid hydrate (BCA; Alfa Aesar, Cat. No. A17491), 6-aminonicotinamide (6-AN; Thermo Fisher Scientific, Cat. No. 104150010), and rotenone (Abcam, Cat. No. ab143145). Compounds were prepared according to the manufacturers’ instructions. Compound final concentrations are provided in the corresponding figure legends.

### Cellular respiration

Cellular respiration was measured using a custom-built gas exchange system as previously described (Eskla *et al*, 2022). The system enables simultaneous real-time measurements of oxygen (O_2_) consumption and carbon dioxide (CO_2_) production in adherent cells. HKC8 cells were seeded at a density of 8 × 10^6^ cells per 100-mm dish on gelatin-coated glass discs and cultured overnight. One hour before the measurements, growth medium was replaced with bicarbonate-free DMEM (Sigma-Aldrich, Cat. No. D5030) supplemented with 5.5 mM D-glucose (Sigma-Aldrich, Cat. No. G7021), 4 mM L-glutamine (Sigma-Aldrich, Cat. No. G7513), and 20 mM HEPES (Sigma-Aldrich, Cat. No. H3375). Carbonic anhydrase (Sigma-Aldrich, Cat. No. C2624) was added immediately before measurements to a final concentration of 25 μg/mL. Cell mass was estimated by optical absorption using an integrating sphere, as previously described (Eskla *et al*., 2022). Optical absorption was determined from the peak at 415–425 nm and expressed relative to the absorption of a calibrated black reference. This method was previously validated against cell number (r^2^ = 0.99). Cells were transferred to the respiration chamber, where O_2_ consumption and CO_2_ production were continuously recorded under the experimental conditions described below:

1. For anoxia-reoxygenation experiments, cells were measured under normoxic conditions (∼1.7% O_2_, 550 ppm CO_2_, balance N_2_), followed by anoxia (O_2_ < 0.03%, 550 ppm CO_2_, balance N_2_) and subsequent reoxygenation under normoxic conditions. In this system, cells are covered by a thin (∼0.1 mm) medium layer, and we have previously shown that ∼1.7% O_2_ in the gas phase is not limiting for cytochrome c oxidase activity and is therefore sufficient to support normoxic respiration (Eskla *et al*, 2022).
2. For hypothermia experiments, cells were initially measured under normoxic conditions at 37 °C, after which the chamber temperature was reduced to 32 °C. Respiration was measured under normoxia at 32 °C, followed by anoxia at 32 °C and subsequent reoxygenation while maintaining the cells at 32 °C. The chamber was then rewarmed to 37 °C under normoxic conditions.
3. For substrate and inhibitor experiments, respiration was first recorded under normoxic baseline conditions in complete respiration medium. Metabolic substrates or pharmacological inhibitors were then added directly to the measurement chamber, followed by normoxic measurements before induction of anoxia. Cells treated with 6-aminonicotinamide were pre-incubated for 16 h before respiration measurements.
4. To assess the contribution of medium acidification to CO_2_ efflux, respiration measurements were performed in media containing 10, 20, or 40 mM HEPES. Medium samples (∼0.6 mL) were collected at three time points: before placing the cells in the measurement chamber (baseline), at the end of the normoxic period, and at the end of a subsequent normoxia–anoxia period. Following the second sampling, cells were rinsed with their original medium to restore baseline pH before the normoxia–anoxia period. Sample pH was measured using a SevenCompact S220 pH meter equipped with an InLab Micro Pro-ISM electrode. The measured pH values and theoretical CO_2_ fluxes attributable to pH-dependent bicarbonate loss are presented in Supplementary Table EV1.

### Differential contribution of glucose and glutamine to CO_2_ using labeled carbon sources

#### Experimental design and ^13^C-labelled substrates

To determine the differential contribution of glucose and glutamine to CO_2_ production, cells were incubated in medium containing glucose and glutamine, with one of the two substrates uniformly ^13^C-labeled (^13^C-glucose, CortecNet, Cat. No. CC860P1; ^13^C-glutamine, Cambridge Isotope Laboratories Inc., Cat. No. CLM-1822-H). Because the LI-COR CO_2_ analyzer detects only approximately 31% of ^13^CO_2_ (McDermitt *et al,* 1995), replacing an unlabeled substrate with its ^13^C-labeled counterpart lowers the measured CO_2_ flux by 69% of the CO_2_ derived from that substrate. The calculation assumes that label substitution does not alter total cellular CO_2_ production.

#### Normalization to respiratory activity

Each experimental setup consisted of individual series for the unlabeled medium (control), ^13^C-glucose, and ^13^C-glutamine, with 3–5 replicates performed for each condition. Absolute respiratory activity varied between experiments, so all series were brought to a common scale using O_2_ consumption, which is not affected by isotope labeling. Uncorrected O_2_ flux differed by approximately 2% between labeled and unlabeled media, whereas differences between experiments were larger (Fig EV1). Assuming that flux time series differ between series mainly by a constant multiplicative factor, one correction factor per series was calculated from the O_2_ fluxes of all series. The transient first three timepoints after chamber closure and after each change of condition (approximately 8 min), during which fluxes stabilized, were excluded. Timepoints during anoxia (3137–4030 s), when O_2_ flux was zero, could not be used for estimating the factors. For each series *i* and each remaining timepoint *t*, a reference value was calculated as the median log-transformed O_2_ flux across all series:

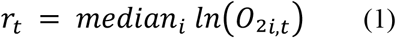

The offset of each series from this reference was calculated as its median over timepoints:

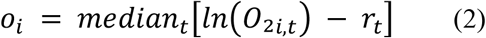

and converted into a multiplicative correction factor:

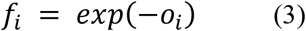

The factors were divided by their geometric mean, so that the final factors had a geometric mean of 1 and the overall flux scale of the dataset was unchanged (*n* = 12):

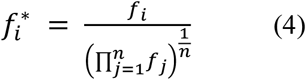

The entire CO_2_ time series of each series was multiplied by the factor derived from its own O_2_ flux:

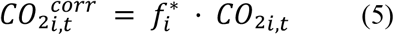

The factors were also applied to the anoxic phase, on the assumption that they reflect differences in the amount of respiring cells between series rather than condition-specific changes. The final factors ranged from 0.91 to 1.07. This step assumes that CO_2_ production scales proportionally with O_2_ consumption between series. Consistent with this, the factors, which were estimated from O_2_ alone, reduced the mean between-series coefficient of variation of O_2_ flux from 6.2% to 2.4% and that of control CO_2_ flux from 6.9% to 3.2%.

#### Substrate contributions

At each timepoint, the CO_2_ flux derived from the labeled substrate S (glucose or glutamine) was calculated as:

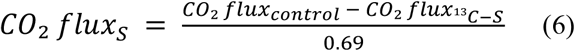

where CO_2_ flux_control_ is the mean corrected CO_2_ flux of the four control series at the same timepoint, CO_2_ flux^13^_C-S_ is the corrected CO_2_ flux of a series in which S was ^13^C-labeled, and 0.69 = 1 ™ 0.31 is the undetected fraction of ^13^CO_2_. Glucose- and glutamine-derived fluxes were calculated independently from their respective labeling series.

#### Residual CO_2_ calculation and interpretation

Residual CO_2_ was calculated by subtracting the estimated glucose- and glutamine-derived contributions from the total CO_2_ flux measured in experiments containing natural-abundance substrates:

*Residual CO_2_ = Total CO_2_ ™ Glucose-derived CO_2_ ™ Glutamine-derived CO_2_*

Residual CO_2_ therefore represents the component of measured CO_2_ flux that could not be attributed to either supplied ^13^C-labelled substrate under the experimental conditions. This component may include contributions from incompletely labelled intracellular glucose and glutamine pools as well as other carbon sources. Because the substrate-derived contributions were estimated independently, small negative residual values could occur when their summed estimates exceeded the corresponding total CO_2_ flux.

### Temperature dependence (Arrhenius) analysis

For Arrhenius analysis, CO_2_ production was measured at 37, 32, 28 and 22 °C under normoxic and anoxic conditions. Apparent activation energies were determined from the Arrhenius equation,

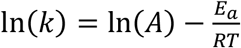

where k is the measured CO_2_ flux, A is the pre-exponential factor, E_a_ is the apparent activation energy, R is the universal gas constant and T is the absolute temperature. ln(k) was plotted against 1/RT, and E_a_ was obtained as the negative slope of the linear regression for each condition.

### Generation of mtDNA-depleted (rho0-like) cells

HepG2 cells were cultured in low-glucose MEM supplemented with 10% fetal bovine serum, 1× penicillin/streptomycin, 0.2 μg/mL ethidium bromide (EtBr), 1 mM sodium pyruvate (Corning, Cat. No. 25-000-CI), and 50 μg/mL uridine (Sigma-Aldrich, Cat. No. U3750) for 96 h to induce mitochondrial DNA depletion. Fresh medium containing the same supplements was replaced after 48 h. Cells were subsequently seeded onto gelatin-coated glass discs for respiration measurements. During respiration experiments, EtBr was omitted from the measurement medium, whereas pyruvate (1 mM) and uridine (50 μg/mL) were maintained as indicated. Untreated HepG2 cells cultured in parallel served as controls.

### Mitochondrial DNA copy number

Mitochondrial DNA depletion was confirmed by determining the mitochondrial-to-nuclear DNA (mtDNA/nDNA) ratio using quantitative PCR, as previously described (Eskla *et al*, 2022). HepG2 cells were seeded at a density of 400,000 cells per well in 6-well plates and cultured overnight. DNA was extracted using 1× PCR buffer (Bioatlas, Cat. No. BA00151) containing proteinase K (final concentration 200 μg/mL) (Thermo Scientific, Cat. No. EO0491). Samples were incubated at 56 °C for 1 h, heated to 97 °C for 20 min to inactivate proteinase K, centrifuged at 14,000 × *g* for 5 min, and stored at ™20 °C until analysis. Quantitative PCR was performed using HOT FIREPol EvaGreen qPCR Supermix (Solis BioDyne, Cat. No. 08-36-00001) on a QuantStudio 12K Flex Real-Time PCR System (Applied Biosystems, Waltham, MA, USA). Mitochondrial DNA was quantified using primers targeting *mt-CytB*, while the single-copy nuclear gene *H4C* served as the reference. Relative mitochondrial DNA content was expressed as the mtDNA/nDNA ratio.

### Statistical analysis

For normally distributed data, comparisons between two groups were performed using an unpaired two-tailed Welch’s t-test. Comparisons involving multiple groups or experimental factors were performed using one- or two-way ANOVA, as appropriate, followed by the multiple-comparisons test specified in the corresponding figure legend. When data did not meet the assumptions of normality, the two-tailed Mann-Whitney test was used. Correlations were assessed by simple linear regression.

## CONCLUSION

Direct simultaneous measurement of CO_2_ and O_2_ fluxes reveals that cellular decarboxylation does not cease when oxygen consumption stops. Anoxic CO_2_ production comprises a substantial glutamine-derived component and a glucose-derived component associated with the pentose phosphate pathway. Pyruvate and α-ketobutyrate stimulate anoxic CO_2_ efflux, with α-ketobutyrate selectively increasing the glucose-derived component, consistent with a role for LDH-dependent cytosolic NAD^+^ regeneration. These findings demonstrate that the coupling between oxygen consumption and CO_2_ production is conditional rather than obligatory and identify redox availability as a determinant of oxygen-independent decarboxylation. Direct CO_2_ measurement therefore reveals metabolic activity that is not captured by oxygen-based respirometry.

## Supporting information

Expanded View

## AUTHOR CONTRIBUTION

H.V. performed experiments, analyzed data, prepared visualizations, and contributed to manuscript revision. H.E. performed experiments, analyzed data, contributed to methodology development, and reviewed the manuscript. U.G. performed experiments and reviewed the manuscript. H.L. contributed to study conceptualization, methodology, data analysis and interpretation, and manuscript revision. K.L.E. conceived and supervised the study, contributed to methodology and data interpretation, administered the project, acquired funding, and wrote the original manuscript with contributions from all authors.

## ACKNOWLEDGEMENTS

We thank Professor Sir Peter J. Ratcliffe for providing the HKC8 cells. This work was supported by Estonian Research Council, PSG959.

## REFERENCES

Arnold PK & Finley LWS (2023) Regulation and function of the mammalian tricarboxylic acid cycle. Journal of Biological Chemistry 299: 102838

Chinopoulos C (2013) Which way does the citric acid cycle turn during hypoxia? The critical role of α-ketoglutarate dehydrogenase complex. Journal of Neuroscience Research 91: 1030–1043

Eskla K-L, Vellama H, Tarve L, Eichelmann H, Jagomäe T, Porosk R, Oja V, Rämma H, Peet N, Laisk A, et al (2022) Hypothermia Alleviates Reductive Stress, a Root Cause of Ischemia Reperfusion Injury. Int J Mol Sci 23: 10108

Grimm F, Asuaje A, Jain A, Silva dos Santos M, Kleinjung J, Nunes PM, Gehrig S, Fets L, Darici S, MacRae JI, et al (2024) Metabolic priming by multiple enzyme systems supports glycolysis, HIF1α stabilisation, and human cancer cell survival in early hypoxia. EMBO J 43: 1545–1569

Hoxhaj G, Ben-Sahra I, Lockwood SE, Timson RC, Byles V, Henning GT, Gao P, Selfors LM, Asara JM & Manning BD (2019) Direct stimulation of NADP+ synthesis through Akt-mediated phosphorylation of NAD kinase. Science 363: 1088–1092

Kory N, Uit de Bos J, van der Rijt S, Jankovic N, Güra M, Arp N, Pena IA, Prakash G, Chan SH, Kunchok T, et al (2020) MCART1/SLC25A51 is required for mitochondrial NAD transport. Sci Adv 6: eabe5310

McDermitt DK, Welles JM & Eckles RD (1995) Effects of Temperature, Pressure and Water Vapor on Gas Phase Infrared Absorption by CO2. LI-COR, Inc.

Messier AA (1975) First apparent dissociation constant of carbonic acid, pK1’, of plasma and erythrocytes. Respiration Physiology 24: 51–68

Mookerjee SA, Goncalves RLS, Gerencser AA, Nicholls DG & Brand MD (2015) The contributions of respiration and glycolysis to extracellular acid production. Biochimica et Biophysica Acta (BBA) - Bioenergetics 1847: 171–181

Ravasz D, Bui D, Nazarian S, Pallag G, Karnok N, Roberts J, Marzullo BP, Tennant DA, Greenwood B, Kitayev A, et al (2024) Residual Complex I activity and amphidirectional Complex II operation support glutamate catabolism through mtSLP in anoxia. Sci Rep 14: 1729

Snell FM (1960) Electrometric measurement of carbon dioxide and bicarbonate ion. J Appl Physiol 15: 729–732

Spinelli JB, Rosen PC, Sprenger H-G, Puszynska AM, Mann JL, Roessler JM, Cangelosi AL, Henne A, Condon KJ, Zhang T, et al (2021) Fumarate is a terminal electron acceptor in the mammalian electron transport chain. Science 374: 1227–1237

Sullivan LB, Gui DY, Hosios AM, Bush LN, Freinkman E & Vander Heiden MG (2015) Supporting Aspartate Biosynthesis Is an Essential Function of Respiration in Proliferating Cells. Cell 162: 552–563

Titov DV, Cracan V, Goodman RP, Peng J, Grabarek Z & Mootha VK (2016) Complementation of mitochondrial electron transport chain by manipulation of the NAD+/NADH ratio. Science 352: 231–235

