## Supplementary material for "Decoupling of CO_2_ and O_2_ fluxes reveals oxygen-independent decarboxylation in anoxic mammalian cells": Expanded View

**Table EV1. Estimated $CO_2$ production from pH-dependent bicarbonate loss at three HEPES concentrations.** The medium was equilibrated with 550 ppm $CO_2$ at 37 °C and treated as an open system; dissolved $CO_2$ was therefore assumed to remain constant during the measurement. Dissolved $CO_2$ was calculated as 12.58 µM using a blood-plasma $CO_2$ solubility coefficient of 22.875 mmol·L$^{-1}$·atm$^{-1}$ (Snell, 1960). Bicarbonate concentration was estimated from pH using the Henderson–Hasselbalch equation $[HCO_3^-] = [CO_2] \times 10^{(pH - pK_a)}$, with $pK_a = 6.1$ (Messier, 1975). Bicarbonate loss during anoxia was calculated relative to the end-of-normoxia pH (see Methods). Bicarbonate loss was converted to nanomoles for the 0.6 mL medium volume on the disc, divided by the measurement period (1,450 s) and normalised to the mean cell number in each condition. Because all bicarbonate lost from the medium was assumed to leave as gas-phase $CO_2$, these values represent a theoretical upper bound. pH values are mean (SD) of three independent experiments.

| HEPES (mM) | Stage | pH (SD) | $HCO_3^-$ (µM) | $\Delta HCO_3^-$ (µM) | $CO_2$ from 0.6 mL (nmol) | $CO_2$ flux (pmol·s$^{-1}$·(10$^6$ cells)$^{-1}$) |
|---|---|---|---|---|---|---|
| 40 | Start | 7.41 (.01) | 256.88 | - | - | - |
| | End of normoxia | 7.30 (.06) | 200.94 | 55.94 | 33.56 | 2.61 |
| | End of anoxia | 7.16 (.08) | 143.35 | 57.59 | 34.55 | 2.69 |
| 20 | Start | 7.36 (.02) | 228.94 | - | - | - |
| | End of normoxia | 7.24 (.01) | 175.01 | 53.93 | 32.36 | 2.45 |
| | End of anoxia | 7.08 (.14) | 119.23 | 55.78 | 33.47 | 2.54 |
| 10 | Start | 7.36 (.04) | 227.19 | - | - | - |
| | End of normoxia | 7.21 (.03) | 160.84 | 66.35 | 39.81 | 2.94 |
| | End of anoxia | 6.72 (.14) | 52.05 | 108.79 | 65.28 | 4.82 |

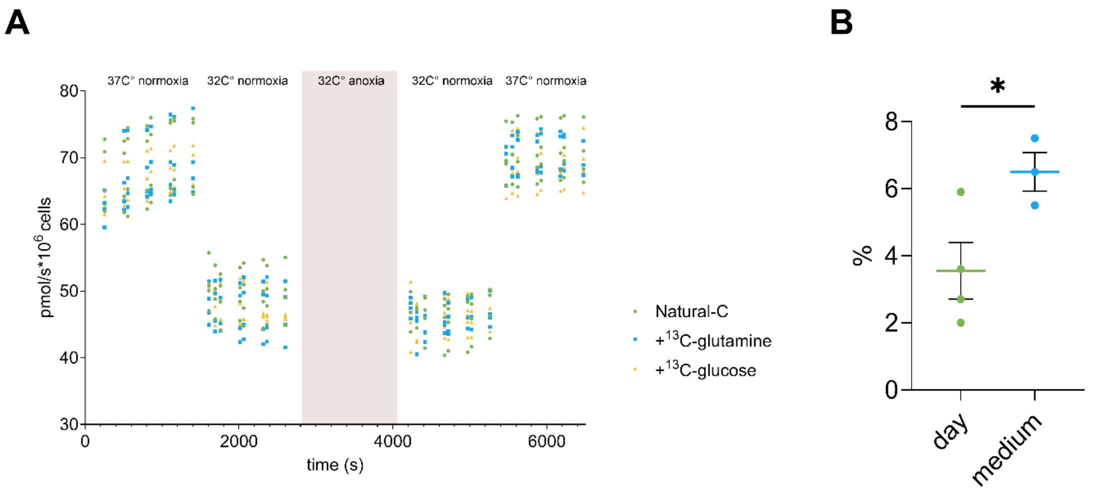


**Figure EV1. Variability in $O_2$ flux measurements across experimental days and substrate conditions. (A)** Oxygen ($O_2$) flux time courses for all HKC8 experiments performed with $^{13}C$-glutamine, $^{13}C$-glucose or unlabeled natural-abundance carbon substrates (Natural-C). **(B)** Mean coefficient of variation (CV) of $O_2$ flux among experiments performed on the same day and among experiments conducted in the same medium type (with $^{13}C$-glutamine, $^{13}C$-glucose, or unlabeled natural-abundance carbon substrates). Data are presented as mean ± SEM. Statistical analysis: (B) two-tailed unpaired Welch's *t*-test. * $P < 0.05$
